# Inferotemporal Cortex Joins the Circuit Before the Code: Non-Serial Inter-Area Synergy in the Macaque Ventral Stream

**DOI:** 10.64898/2026.05.06.723331

**Authors:** Arun Ram Ponnambalam, Krishnan Venkiteswaran Pottore

## Abstract

The ventral visual stream is widely modeled as a serial feedforward hierarchy in which V1, V4, and IT population codes develop sequentially during object recognition. We ask whether a second, concurrent coding mode exists—one organized not by anatomical order but by joint population structure across areas. Using Partial Information Decomposition applied to simultaneous multielectrode spiking recordings across all three areas at millisecond resolution—the first simultaneous three-area spiking PID analysis of the primate ventral stream—in two macaque monkeys viewing 25,000+ natural images, we decompose population coding into serial (unique per area) and synergistic (joint across areas) components at 5 ms resolution across five CNN target representations spanning low-level spatial features to high-level object identity. Three findings replicate across both animals and all five representations. First, synergistic inter-area coupling emerges before IT carries any unique object-related information—a dissociation of 15–65 ms that replicates in direction without exception across both animals—such that the joint population integrates before the apex encodes; moreover, V1–IT synergy persists for over 120 ms after V1’s unique information reaches zero. Second, although V1↔IT and V1↔V4 coupling emerge simultaneously and rise in parallel, V1↔IT exhibits stronger peak synergy at mid-to-high-level targets in both animals, suggesting a dominant role for non-serial joint coding. Third, when V1 and V4 are treated as an integrated feedforward block, their synergistic coupling with IT emerges last across all tested conditions—the feedforward foundation is the final component to join the synergistic mode, not the first. Together, these results show that serial and synergistic population codes co-occur in the same recordings, overlap in time, but follow different organizational principles, Providing a new level of nuance in our understanding of the primate ventral stream and introducing concrete constraints for biologically grounded models of vision.

## 1 Introduction

The primate ventral visual stream maps a continuously varying retinal image onto a stable, transformation-tolerant representation of object identity within roughly 200 ms. The canonical account holds that this is achieved through a serial feedforward hierarchy in which V1 encodes oriented edges and spatial frequencies, V4 integrates mid-level features such as curvature, colour, and texture, and inferotemporal cortex (IT) arrives at a transformation-tolerant representation of object identity [1]. This cascade has been operationalised by deep CNNs trained on image classification: task-optimised hierarchical networks predict single-unit responses in macaque V4 and IT [3], earlier layers best predict V1 and V2 while deeper layers best predict V4 and downstream regions [5], and match quality increases with model depth and categorisation performance [2]. Deep supervised networks far exceed all previously tested model classes—including HMAX, VisNet, and hand-crafted feature models—in explaining IT representational geometry [4] and predicting individual IT spiking responses [3].

A strictly serial cascade carries specific, falsifiable predictions: (i) representational content should emerge in the order V1 → V4 → IT; (ii) inter-area coupling should respect anatomical adjacency, with adjacent pairs (V1–V4, V4–IT) interacting more tightly than the non-adjacent pair (V1–IT); (iii) each area’s unique content should emerge before, not after, it begins to interact with other areas; and (iv) the feedforward foundation V1+V4 should be the first composite stage to couple functionally with IT. Critically, a purely feedforward model produces zero synergy by construction—synergy requires joint information exceeding what any individual area provides, which a deterministic serial cascade cannot generate. Testing whether synergistic coupling exists, when it emerges relative to unique information, and which pairs generate the most synergy therefore provides a direct test of the feedforward account.

Evidence already shows purely feedforward models fail in specific domains. Anatomical mapping reveals inter-areal projection weights spanning five orders of magnitude [11]; pharmacological and optogenetic inactivation of feedback alters response gain, surround suppression, and tuning in downstream regions [10]; and ConvRNNs incorporating recurrence and long-range feedback better match the temporal dynamics of V4 and IT than feedforward-only architectures [7].

Partial Information Decomposition (PID) [9] provides exactly the required framework, decomposing total mutual information into non-negative atoms of unique, redundant, and synergistic information. Large-scale decomposition of human fMRI reveals redundant interactions in sensorimotor cortex and synergistic interactions in higher-order association cortex, with synergy exhibiting the greatest evolutionary expansion from macaque to human [12]. Extended to invasive electrophysiology, inter-areal ECoG in awake marmosets is synergistic across the auditory hierarchy, requiring strong long-range feedback in a brain-constrained model [13]; and in macaque and human visual cortex, broadband V1–V4 signals are highly synergistic while narrowband gamma is predominantly redundant—dissociating the information-theoretic roles of two concurrently active signal types [14].

The THINGS Ventral Stream Spiking Dataset (TVSD) [15]—chronic multi-unit recordings spanning V1, V4, and IT in two macaque monkeys viewing *>*25,000 natural images from the THINGS database [18]—provides the stimulus diversity, temporal resolution, and simultaneous multi-area coverage required to address this question. Population decoding establishes that small IT ensembles carry surprisingly accurate object identity information [16], but decodability from a single area is fundamentally different from asking what the circuit computes collectively. Correlations within and across areas structure both encoding and downstream transmission [17], and prior TVSD characterisations show noise correlations are strongest between similarly tuned neurons [15]—consistent with shared feedforward drive—but whether joint activity carries information beyond individual areas, and when, has not been measured.

Here we apply three-source PID to simultaneous spiking recordings from V1, V4, and IT in both animals viewing *>*22,000 natural images, decomposing unique, redundant, and synergistic information at 5 ms resolution across the 300 ms post-stimulus window for five ResNet-18 targets spanning early convolutional to final identity layers. We report four findings replicating across both animals, all five CNN targets, and five window-stride configurations. First, inter-area synergy emerges 40–65 ms before IT carries any unique object-related information—IT participates as a synergistic circuit partner before it contributes as an independent encoder. Second, pairwise synergy emerges near-simultaneously across areas, with V1–IT and V1–V4 showing concurrent onset and parallel rise. Contrary to anatomical expectations, the non-adjacent V1–IT pair is not weaker but instead matches or exceeds adjacent pairs, indicating a non-serial, distributed mode of inter-areal coding. Third, the V1+V4 feedforward block synergises with IT later than every other block-area combination—the feedforward foundation is last, not first, at 167.5 ms in both animals. Together these findings reveal a ventral stream that simultaneously operates as a serial representational hierarchy—unique information emerging in the V1 → V4 → IT sequence predicted by feedforward models—and as a distributed integrated system in which non-adjacent, non-serial interactions dominate the synergistic code, while the feedforward foundation is not recruited preferentially into the integrated circuit—a pattern not predicted by feedforward models.

## 2 Methods

### 2.1 Dataset and Preprocessing

We analysed the THINGS Ventral Stream Spiking Dataset (TVSD) [15], which provides simultaneous high-channel-count multi-unit activity (MUA) recordings from macaque V1, V4, and IT during passive viewing of 25,248 natural images from the THINGS database [18]. Each trial was recorded over a 300 ms window at 1 ms resolution, with stimulus onset at 100 ms, yielding a 100 ms prestimulus baseline and a 200 ms post-stimulus analysis window. Data from two animals are analysed independently; all main-text findings replicate in both. Monkey F: 512 V1, 320 IT, 192 V4 channels. Monkey N: 960 channels after exclusion of one unstable array (Array 6, 64 channels): 448 V1, 256 V4, 256 IT. All remaining channels were retained. Each channel was z-scored across all time bins and trials jointly to equalise firing-rate scale across electrodes and animals. Neural population activity was summarised in non-overlapping 15 ms windows stepped by 5 ms, yielding 57 time bins with centres spanning 7.5–292.5 ms relative to stimulus onset. For each time bin [*t, t* + *W*) with *W* = 15 ms, the neural observation for trial *k* and ROI *r* was obtained by averaging the z-scored MUA across the window.

### 2.2 CNN Representational Targets

We extracted population-code targets from five stages of a pre-trained ResNet-18 [19]: outputs of residual blocks layer1–layer4 (global-average-pooled to 64, 128, 256, 512 dimensions respectively) and the final avgpool layer (512-d). Each was compressed to 50 PCs via standard PCA, yielding a target matrix **Y**^(*ℓ*)^ ∈ ℝ^*N ×*50^ per layer.

### 2.3 Marchenko–Pastur PCA for Signal Subspace Estimation

For each ROI *r* and time bin *t*, column-centred neural data 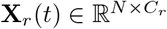 were projected onto a signal subspace estimated via the Marchenko–Pastur law [20, 21]. Noise variance was estimated as the median positive eigenvalue of the sample covariance 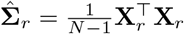,

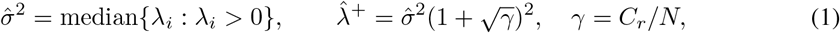

and the signal dimensionality and projection were

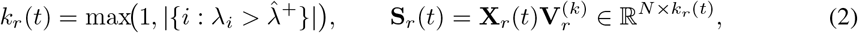

where 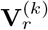 contains the top *k*_r_ (*t*) eigenvectors of 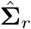; a floor of one component is retained to avoid degenerate representations at bins with no detectable signal. In our regime (*γ* ∈ [0.002, 0.10]; V1: *C*=448, V4/IT: *C*=256, *N*≥22,000), signal components occupy at most 10–30% of the eigenspectrum, so the median falls within the MP noise bulk [21]; any upward bias in 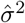 raises 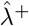 and under-retains signal, a conservative error that can only underestimate synergy and unique information (Section 2.4).

Joint representations for synergy estimation were obtained by concatenating per-ROI projections {**S**_*r*_(*t*)} column-wise and applying a second-level MP-PCA pass to the combined matrix, using the same threshold procedure with updated aspect ratio *γ*_joint_ = ∑_*r*_ *k*_*r*_(*t*)*/N*. This step removes residual noise components that survive single-area filtering and balances contributions from areas with differing electrode counts and intrinsic dimensionalities, so that the resulting joint projection **Z**(*t*) captures inter-area shared structure rather than single-area dominance. A dedicated bias-correction term was estimated separately for **Z**(*t*) (Section 2.4).

### 2.4 Mutual Information Estimation under a Gaussian Model

Let 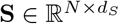 and 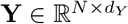 be zero-mean neural and target matrices. Under a jointly Gaussian assumption, the mutual information admits the closed-form log-determinant expression

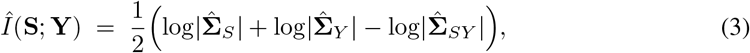

where 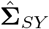 is the joint sample covariance of [**S, Y**] and log-determinants are computed from signed LU factorisation; the estimate is clipped to zero if any determinant sign is non-positive.

The Gaussian plug-in estimator was chosen on theoretical and practical grounds. Under a jointly Gaussian model, mutual information depends only on second-order statistics and thus quantifies the information accessible to optimal linear decoders—an operational proxy for inter-area communication under biologically plausible readouts [30–33]. Accordingly, it measures linear-accessible (secondorder) information rather than total information; higher-order dependencies not reflected in covariance are not captured by design. This aligns with MP-PCA, which emphasizes shared variance across electrodes. Practically, the Gaussian framework remains well-conditioned and tractable in the 50–160D, large-scale regime considered (57 time bins *×* 5 CNN layers *×* 500 shuffles), where *k*-NN and neural estimators are statistically unstable and computationally prohibitive [22]. Robustness was assessed using a Gaussian Copula Mutual Information (GCMI) estimator on a 3,000-sample subset: while absolute magnitudes differed, temporal profiles and ordinal relationships claimed were preserved (corr. ≈ 0.75 synergy; ≈ 0.9 redundancy/unique), indicating estimator-invariant qualitative structure. Pre-stimulus MI was ≈ 0 after shuffle correction, confirming effective bias removal. Together, this supports the Gaussian plug-in as a scalable, empirically validated method for characterizing dominant, linearly accessible inter-area interactions.

#### Shuffle-based bias correction

Finite-sample bias was corrected by subtracting a shuffle-based empirical mean: for each time bin, **Y** was row-permuted *B* = 500 times on a subsample of *N*_sub_ = 5,000 trials, and the bias estimate 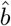 was the mean MI across permutations. Separate bias terms were estimated for each individual-ROI MI term and for each joint representation; for pairwise analyses, the per-area marginal bias terms were taken from the three-source estimation (the marginal MI of a single area is independent of the other sources present), while a dedicated joint bias was estimated for each pair’s joint projection. All bias-corrected estimates satisfy 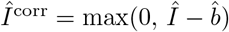. All PID time series were additionally baseline-corrected by subtracting the pre-stimulus mean (bins centred at *<* 100 ms); the same column-wise subtraction was applied independently to each shuffle replicate before any threshold computation, ensuring observed and null values are compared on a consistent change-from-baseline scale.

### 2.5 Partial Information Decomposition

Given neural source representations 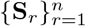 and target **Y**, Partial Information Decomposition [24] decomposes the total joint information into non-negative atoms of redundancy, unique information, and synergy.

#### Redundancy and unique information

We adopt the *I*_min_ redundancy measure of Barrett [23], which for jointly Gaussian variables is the *unique* non-negative decomposition consistent with the axiom that redundancy depends only on the individual source–target marginals [23, 22]; the conceptual concerns with *I*_min_ for discrete variables therefore do not apply in our Gaussian setting. Redundancy is defined as the minimum information any single source carries about the target:

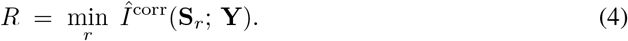

The unique information of source *r* is the excess above this redundant floor:

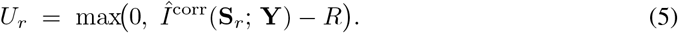

#### Synergistic information

Synergy is the information present in the joint representation that cannot be accounted for by the redundant and unique components:

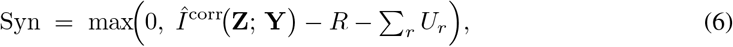

where **Z** is the second-level MP-PCA projection of the concatenated source representations (Section 2.3). The max(0, ·) operations enforce non-negativity of all atoms, following the lattice construction of Williams and Beer [24] applied to the Gaussian framework of Barrett [23]. Because the joint bias 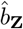 is estimated separately from the marginal biases 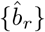, the decomposition avoids the systematic over-correction of the synergy atom that arises from applying a single pooled bias term.

All primary conclusions are drawn from ordinal and temporal comparisons—the relative ordering of synergy across area pairs, block configurations, and CNN target layers, and the timing of synergy onset relative to unique IT information emergence—which are robust to any monotone rescaling of MI values. This insulates the key findings from any residual dependence of absolute MI magnitude on distributional assumptions.

#### Three-source hybrid PID

For the three-area (V1, V4, IT) decomposition, sources were **S**_V1_, **S**_V4_, **S**_IT_. Redundancy was *R* = min_*r*_ *Î*^corr^(**S**_*r*_; **Y**), unique information was computed per area, and synergy was computed against the joint second-level projection **Z**_all_ of all three sources.

#### Pairwise PID

For each of the three area pairs (*r*_*A*_, *r*_*B*_) ∈ {(V1, IT), (V1, V4), (IT, V4)}, an independent two-source PID was computed using 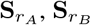, and their dedicated second-level joint projection 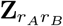.

#### Block PID

To directly test the temporal ordering of feedforward integration—specifically whether the V1+V4 block, representing the canonical feedforward input to IT, synergises with IT before or after other area combinations—pairs of areas were treated as a single composite source. For each of the three block configurations— {(V1 +IT vs. V4), (V1 +V4 vs. IT), (IT+V4 vs. V1)} —the pair block was represented by the second-level MP-PCA projection of its two constituent areas’ individual signal projections, following the same procedure as the three-source joint representation described above. A two-source PID was then computed between this block projection and the singleton area’s signal projection.

### 2.6 Statistical, Representational, and Robustness Analyses

Two complementary permutation tests were applied to each scalar time series (Total MI, Synergy, and pairwise/block synergy components): an **omnibus test** comparing the mean post-stimulus (≥ 100 ms) value to *B*=500 shuffle means to obtain a single *p*-value, and a **cluster-mass test** that identified contiguous above-threshold bins (threshold: *µ*_null_(*t*) + 2*σ*_null_(*t*)), summed their values, and assessed significance at *α*=0.05 against the maximum-cluster-mass null distribution. Cluster onset is defined as the first bin of the first significant cluster and provides an approximate lower bound on effect onset; the cluster-mass test does not license precise latency inference [28]. Representational similarity analysis (RSA) [27] was performed between each area’s MP-PCA projections and CNN targets at each time bin using Spearman correlation of pairwise Euclidean-distance RDMs on 500-trial subsamples (5 repetitions, mean taken), with significance assessed via 200 label permutations at *α*=0.05; the earliest significant bin defined representational onset for each area and served as the reference for comparing synergy onset. Robustness was evaluated by repeating the three-area hybrid PID across five window/step configurations (*W, s*) ∈ {(10, 2), (15, 4), (20, 5), (30, 5), (50, 10)} ms using 5,000 trials each, confirming that synergy onset and peak magnitude—and thus all qualitative conclusions—remained consistent across the full parameter sweep (Supplementary Fig. S1).

## Results

We recorded simultaneous MUA across V1, V4, and IT in two macaque monkeys (Monkey F: 509 V1, 311 IT, 192 V4 electrodes; Monkey N: 448 V1, 256 IT, 256 V4 electrodes; 25,248 trials each) passively viewing 22,248+ natural images from the THINGS dataset [15]. We applied Partial Information Decomposition (PID) with per-area Marchenko–Pastur PCA (MP-PCA) noise rejection at 5 ms temporal resolution, targeting five ResNet-18 representations from low-level spatial features (Layer 1, 64-d) to high-level object identity (AvgPool, 512-d). Pre-stimulus MI is exactly 0.000000 nats in all conditions, confirming successful bias correction; significance was assessed via omnibus and cluster-mass permutation tests (500 label-shuffled shuffles). All qualitative conclusions the synergy-precedes-IT-unique gap (15–65 ms), pairwise ordering at the object-identity target, and block configuration ordering — are invariant across five window–stride configurations (10–50 ms windows; Supplementary Figure S1).

### 3.1 Sequential feedforward sweep

As shown in 1, RSA onset and PR quench both follow V1 → V4 → IT in both animals (RSA onset, Monkey F: 122.5, 137.5, 152.5 ms; Monkey N: 127.5, 132.5, 137.5 ms; PR quench percentages and absolute timings in Supplementary Table S1), confirming the feedforward component is present and providing the reference timeline against which synergy onset is compared below.

**Figure 1:**
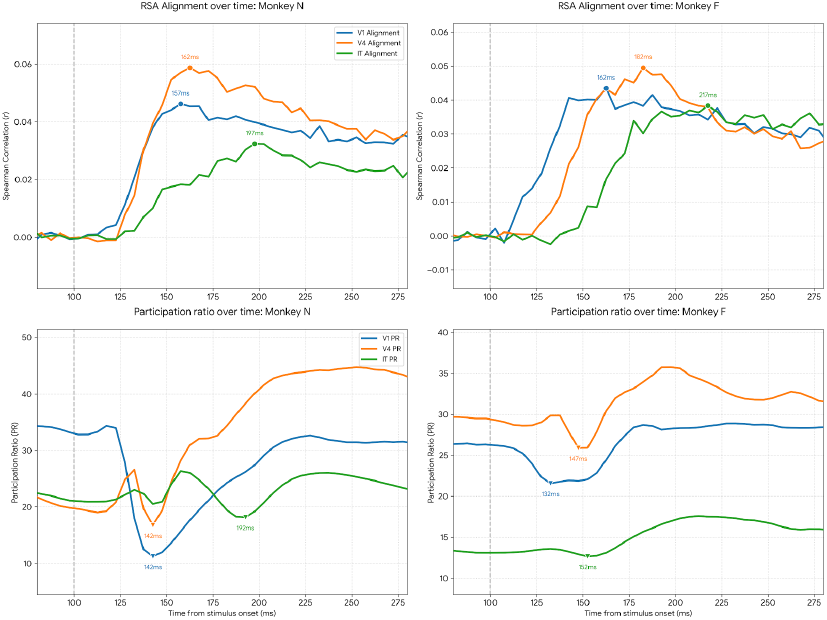
Sequential feedforward sweep in both animals. *Top row:* RSA alignment (Spearman *r*) between each area’s MP-PCA projections and AvgPool CNN features. Onset follows V1 → V4 → IT in both animals. *Bottom row:* Participation ratio (PR) quench timing follows the same hierarchy. All timings reported relative to stimulus onset (dashed line).

### 3.2 Joint population code: magnitude and robustness

Total MI between the joint V1+V4+IT population and CNN features is large, statistically robust, and consistent across animals. At AvgPool — the object-identity target and the tightest cross-animal comparison — peak total MI reaches 1.594 nats (Monkey F) and 1.587 nats (Monkey N), a 0.4% magnitude difference and a 5 ms timing difference, the closest replication in the dataset. Per-layer MI values across all five CNN targets are provided in Supplementary Table S2; briefly, total MI peaks at mid-level CNN layers (Layer 3: 3.365 nats Monkey F, 3.371 nats Monkey N) and falls at Layer 4 and AvgPool because global average pooling discards the spatial structure that dominates mid-level representations. Cluster-mass tests yield *p*=0.002 and omnibus tests yield *p*=0.002 at all five layers in both animals; not one of 500 shuffles exceeded the observed signal at any layer in either animal. Temporally, total MI at AvgPool rises steeply from onset (∼142.5 ms Monkey F; ∼137.5 ms Monkey N), peaks within 45–55 ms of onset, and remains elevated throughout the 300 ms window.

### 3.3 inter-area synergy begins before IT carries unique object information

The serial feedforward account predicts that individual area representations should precede inter-area coupling. We find the opposite for IT, consistently across all layers and both animals.

Synergy — information accessible only from the joint population and absent from any single area — is statistically significant at all layers in both animals (cluster-mass *p*=0.002, omnibus *p*=0.002 throughout). It emerges substantially before IT unique information at every measurable layer. At AvgPool specifically: synergy onset (first significant cluster bin) is 137.5 ms (Monkey F) and 132.5 ms (Monkey N), while IT unique information onset is 167.5 ms (Monkey F) and 177.5 ms (Monkey N) — gaps of 30 ms and 45 ms respectively. Across all measurable layers the gap ranges from 15–65 ms: smallest at Layers 1–2 in Monkey F (15 ms), largest at Layer 3 in Monkey N (65 ms), with the gap consistently smaller at higher CNN layers in both animals 2 panel C; full per-layer onset timings are given in Supplementary Table S3. IT unique information is absent in Monkey N for Layers 1–2, consistent with Monkey N having fewer IT electrodes (256 vs. 311 in Monkey F) and correspondingly lower sensitivity to features IT does not primarily encode.

As shown in 2 panel A, Synergy magnitude peaks at 1.562 nats in Monkey F and 1.343 nats in Monkey N at Layer 3, and at 0.712 nats and 0.603 nats at AvgPool. At AvgPool, synergy rises sharply to its peak within the first 40 ms post-onset and then decays to a sustained plateau — remaining above 0.15 nats from 162.5 ms to the end of the window in Monkey F and above 0.44 nats from 172.5 ms onwards in Monkey N — demonstrating that inter-area synergistic coding is not a transient initialisation event but a persistent feature of the joint population response. MP-PCA confirms that all three areas carry substantial independent signal throughout: at AvgPool in Monkey F, V1 retains ≈162 signal PCs pre-stimulus (145 on average post-stimulus), IT≈115, and V4≈69, ruling out noise dominance in any area. V1–IT pairwise synergy sustains at 0.244 nats for over 120 ms *after* V1’s unique information reaches zero (at 167.5 ms) while IT’s unique representation simultaneously reaches its maximum (1.018 nats at 237.5 ms, Monkey F). Synergy does not vanish once the representational handoff is complete — it persists when both areas are in stable post-transition states.As shown in 2 panel D, synergy peaks in Layer 3—reaching approximately 1.56 bits in Monkey F and 1.34 bits in Monkey N—before dropping by roughly 50% in Layer 4 and the AvgPool stage (0.71 and 0.60 bits, respectively). This suggests that Layer 3 acts as a mid-level scaffold where multi-level synergistic coding across different ROIs is maximally correlated.

**Figure 2:**
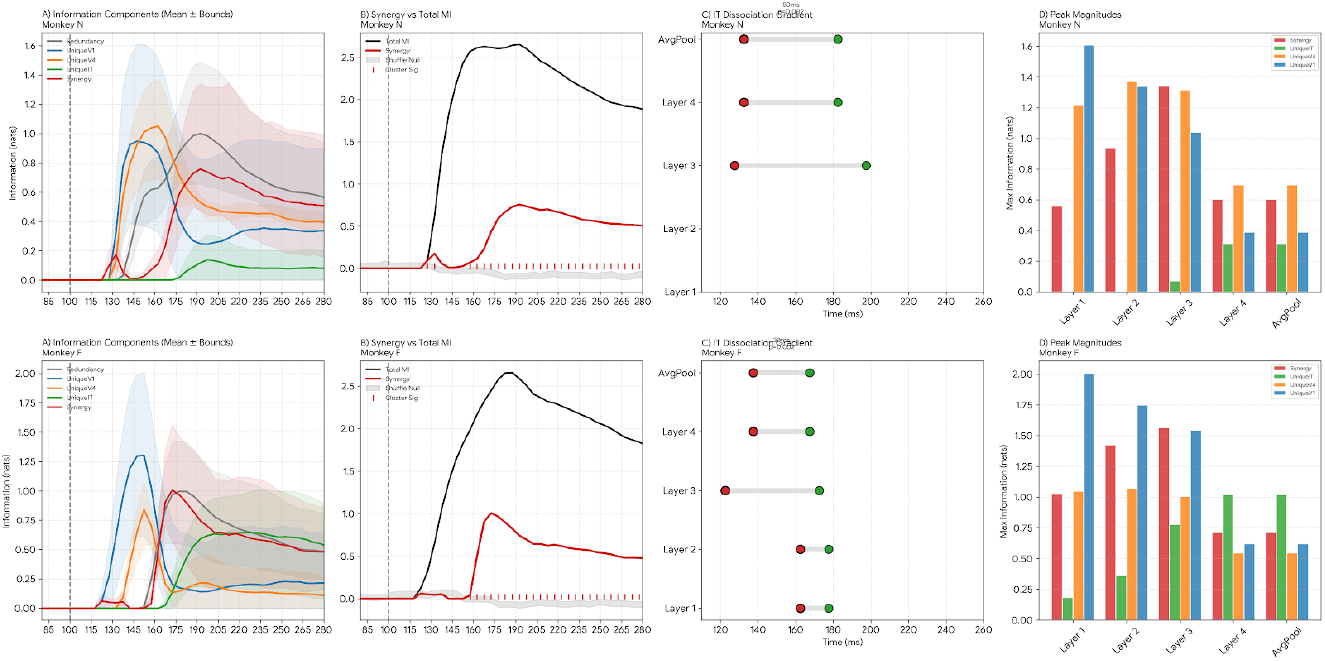
Synergy precedes IT unique information across all layers and both animals. *Left:* Full PID decomposition (Layer 4) showing synergy (red) emerging before the IT unique component (green). *Middle:* Synergy (red dashed) and total MI (black) at AvgPool with shuffle null (grey); significant bins marked. *Right:* IT Dissociation Gradient — red dot marks synergy onset, green dot marks IT unique onset, per layer per animal. Gap ranges from 15–65 ms with direction replicating without exception.

**Figure 3:**
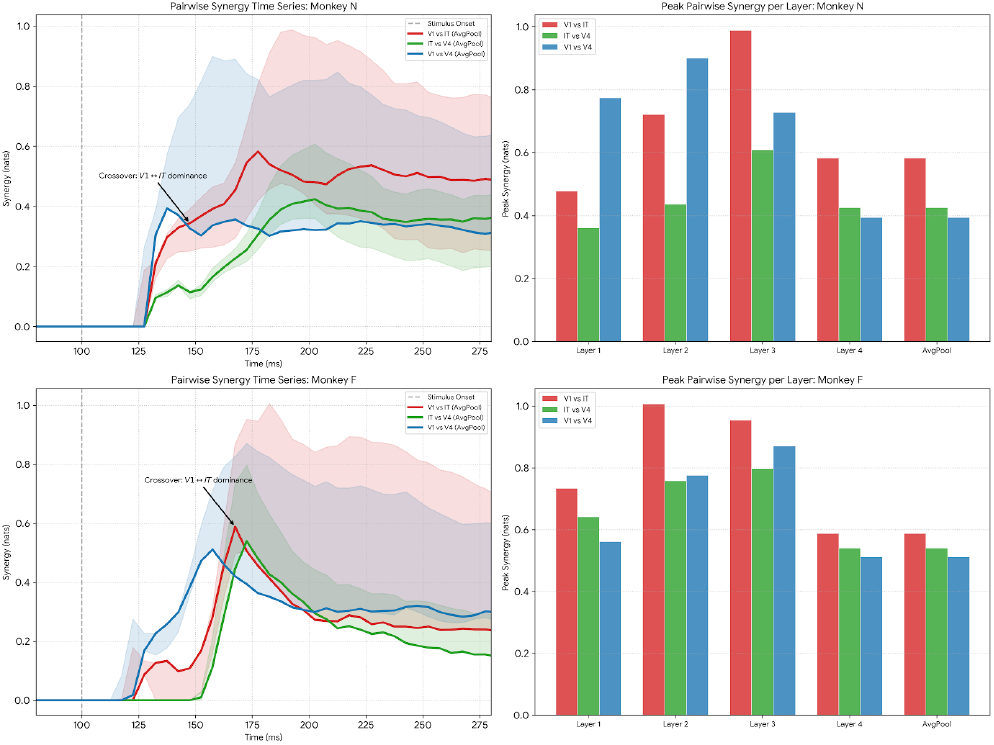
Pairwise synergy emerges non sequentially. *Left:* Pairwise synergy time series (Layer 4) for each monkey. V1↔IT (red) and V1↔V4 (blue) rise in parallel before V1↔IT takes the lead at 167.5 ms (Monkey F, annotated arrow). *Right:* Peak pairwise synergy across all five CNN layers. V1↔IT is strongest at all layers in Monkey F and from Layer 3 onward in Monkey N.

### 3.4 Near-Parallel, Non-Serial Onset of Inter-Areal Synergistic Coupling

Anatomical adjacency predicts that directly adjacent pairs, specifically V1–V4 and IT–V4, and then recurrent feedback, plausibly V1–IT in that order, should proceed sequentially in terms of pairwise synergistic coupling. However, we observe that the pairwise synergies V1–IT and V1–V4 begin simultaneously and also start their ascent near-simultaneously. V1–IT and V1–V4 share the same feedforward onset timing—0 ms apart in Monkey N (all pairs at 132.5 ms) and 5 ms apart in Monkey F (V1–V4 at 122.5 ms, V1–IT at 127.5 ms)—and rise in parallel throughout the steep post-onset phase. This suggests parallel collaborative coding of the image representation. It is also expected that adjacent pairs generate more synergy than the non-adjacent pair V1–IT. This prediction is falsified at the object-identity target in both animals. At the AvgPool layer, the non-adjacent pair is the strongest in both subjects: Monkey F: V1–IT (0.587 nats) *>* IT–V4 (0.539 nats) *>* V1–V4 (0.511 nats). Monkey N: V1–IT (0.582 nats) *>* IT–V4 (0.424 nats) *>* V1–V4 (0.393 nats). This ordering holds across Layers 3,4 and AvgPool in both animals and across all five layers in Monkey F, as shown in 3. At mid-level targets, V1–IT peaks at 1.007 nats (Monkey F, Layer 2), 0.953 nats (Monkey F, Layer 3), and 0.988 nats (Monkey N, Layer 3), remaining the dominant pair from Layer 3 onward in Monkey N. A conservative estimate still finds V1–IT coupling to be nearly as strong as the feedforward pair nearly everywhere. While the lowest layers in Monkey N initially reflect stronger spatial feature coupling (V1–V4 at 0.774/0.900 nats), consistent with low-level spatial encoding, the non-adjacent V1–IT synergy eventually overtakes these adjacent pairs. The hierarchy is determined at the peak magnitude, not the onset: V1–IT overtakes V1–V4 at 167.5 ms in Monkey F and is dominant from the first significant rise in Monkey N, implicating representational content rather than connection timing as the organizing principle. All three pairs emerge near-concurrently: the onset spread at AvgPool is only 30 ms in Monkey F (122.5–152.5 ms) and 0 ms in Monkey N (all three pairs at 132.5 ms), ruling out a serial inter-area synergy dynamic.

**Figure 4:**
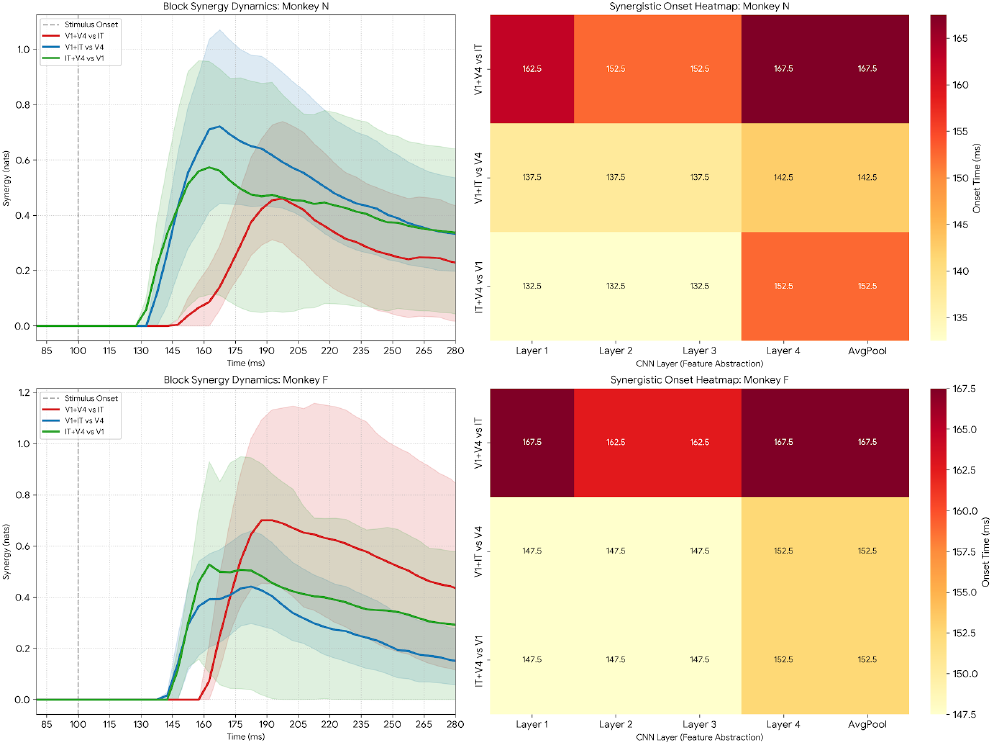
The feedforward foundation synergises with IT last in all ten conditions. *Left:* Block synergy time series (Layer 4). V1+V4 vs. IT (red) rises last. *Right:* Onset heatmap across block configurations and CNN layers — V1+V4 vs. IT row is darkest (latest) in every cell, zero exceptions.

### 3.5 The feedforward foundation synergises with IT last, not first

The feedforward account predicts that V1+V4 — the accumulated feedforward computation — should synergise with IT earliest, since IT is thought to read out the feedforward foundation. We find the opposite in every single condition across both animals and all five CNN layers. V1+V4 vs. IT block synergy always emerges last. In Monkey F, the onset ordering at Layers 1–3 is V1+IT (142.5 ms) → IT+V4 (147.5 ms) → V1+V4 (162.5 ms), and at Layer 4 and AvgPool it is V1+IT (152.5 ms) = IT+V4 (152.5 ms) → V1+V4 (167.5 ms). In Monkey N, the ordering at Layers 1–3 is IT+V4 (132.5 ms) → V1+IT (137.5 ms) → V1+V4 (152.5 ms at Layers 1–2; 147.5 ms at Layer 3), and at Layer 4 and AvgPool it is V1+IT (142.5 ms) → IT+V4 (147.5 ms) → V1+V4 (167.5 ms). V1+V4 vs. IT is latest in all 10 conditions without exception, with a consistent temporal disadvantage of 10–20 ms (2–4 time bins) — well above the 5 ms bin resolution As shown in 4. The animal-level timing offset at high layers (167.5 ms in both animals at AvgPool) is the tightest cross-animal replication in the block analysis.

## 3 Discussion and conclusion

Our results identify two concurrent and complementary computational regimes in the primate ventral stream during object recognition. First, a serial feedforward progression is preserved: unique information and population geometry emerge across V1, V4, and IT in anatomical order, consistent with canonical hierarchical models. Second, and critically, a distributed synergistic code operates in parallel. This code is not detectable from single-area analyses and is not predicted by purely feedforward accounts, but instead reflects inter-area interactions in which regions jointly encode information that is not present in any area alone while maintaining distinct marginal representations. This joint code recruits IT into an inter-area circuit prior to the emergence of stable IT-unique information, and incorporates the feedforward V1+V4 foundation only as the final component to couple with IT. Quantifying its onset timing, inter-area ordering, and relationship to individual-area representations yields explicit constraints on mechanism: non-serial interactions precede the completion of local computations, and joint representations coexist with, rather than replace, area-specific encoding. V1–IT synergy remains substantial 120,ms after both areas reach peak individual information, demonstrating that the circuit encodes information unavailable to either area in isolation. The pairwise structure further constrains the mechanism: V1↔ITV1↔IT V1↔IT emerges at response onset and generates comparable or greater synergy despite being the non-adjacent pair. Any candidate model must therefore account not only for the temporal dynamics of individual areas, but also for the non-serial, concurrent inter-area synergistic coding that underlies object representation in the ventral stream.

### Limitations

We use a Gaussian plug-in MI estimator applied to MP-PCA projections, which captures second-order covariance structure and may not recover higher-order dependencies; accordingly, all reported synergy should be interpreted as jointly decodable structure accessible to linear readouts, rather than total information. Crucially, the temporal precedence and ordering of this structure impose constraints on any model of inter-area communication. In addition, our partial information decomposition relies on *I*_min_, which is known to allocate relatively more mass to synergy compared to alternative definitions such as *I*_dep_ or BROJA. As such, the absolute magnitude of the reported synergy should be interpreted cautiously. However, this bias is not expected to invert the qualitative structure of the results: as established in the Methods, Gaussian copula MI (GCMI) preserves ordinal relationships under monotone transformations, and our central findings concern relative ordering and temporal structure—namely, which area pairs exhibit stronger synergy, the non-adjacent dominance of *V* 1 ↔ *IT*, and the late engagement of the *V* 1 + *V* 4 feedforward block. These features depend on consistent rank and timing relationships rather than absolute values, and are therefore robust to monotone rescaling and moderate estimator bias. The target representations are drawn from a fixed pretrained ResNet-18; different architectures or task-optimized models may yield quantitatively different MI values, although the ordinal structure of the results is expected to generalize. Finally, passive viewing does not systematically engage top-down attentional modulation; whether the synergistic structure we observe is amplified, redistributed, or remains stable under task demands remains an open question.

## 4 Data Availability

The neural population dataset analyzed in this work is derived from previously published research and is publicly accessible via *Neuron* (2024; DOI: 10.1016/j.neuron.2024.00881). To maintain doubleblind anonymity during the review process, we cite this dataset in the third person. Instructions for integrating the dataset with our provided code can be found in the accompanying README.md.

## NeurIPS Paper Checklist

The checklist is designed to encourage best practices for responsible machine learning research, addressing issues of reproducibility, transparency, research ethics, and societal impact. Do not remove the checklist: **The papers not including the checklist will be desk rejected**. The checklist should follow the references and follow the (optional) supplemental material. The checklist does NOT count towards the page limit.

1. **Claims** Question: Do the main claims made in the abstract and introduction accurately reflect the paper’s contributions and scope? Answer: [Yes] Justification: To the best of our knowledge, claims are thoroughly segmented and elaborated in the results Guidelines:
  - The answer [N/A] means that the abstract and introduction do not include the claims made in the paper.
  - The abstract and/or introduction should clearly state the claims made, including the contributions made in the paper and important assumptions and limitations. A [No] or [N/A] answer to this question will not be perceived well by the reviewers.
  - The claims made should match theoretical and experimental results, and reflect how much the results can be expected to generalize to other settings.
  - It is fine to include aspirational goals as motivation as long as it is clear that these goals are not attained by the paper.
2. **Limitations** Question: Does the paper discuss the limitations of the work performed by the authors? Answer: [Yes] Justification: Please find the limitations section in the main paper. Guidelines:
  - The answer [N/A] means that the paper has no limitation while the answer [No] means that the paper has limitations, but those are not discussed in the paper.
  - The authors are encouraged to create a separate “Limitations” section in their paper.
  - The paper should point out any strong assumptions and how robust the results are to violations of these assumptions (e.g., independence assumptions, noiseless settings, model well-specification, asymptotic approximations only holding locally). The authors should reflect on how these assumptions might be violated in practice and what the implications would be.
  - The authors should reflect on the scope of the claims made, e.g., if the approach was only tested on a few datasets or with a few runs. In general, empirical results often depend on implicit assumptions, which should be articulated.
  - The authors should reflect on the factors that influence the performance of the approach. For example, a facial recognition algorithm may perform poorly when image resolution is low or images are taken in low lighting. Or a speech-to-text system might not be used reliably to provide closed captions for online lectures because it fails to handle technical jargon.
  - The authors should discuss the computational efficiency of the proposed algorithms and how they scale with dataset size.
  - If applicable, the authors should discuss possible limitations of their approach to address problems of privacy and fairness.
  - While the authors might fear that complete honesty about limitations might be used by reviewers as grounds for rejection, a worse outcome might be that reviewers discover limitations that aren’t acknowledged in the paper. The authors should use their best judgment and recognize that individual actions in favor of transparency play an important role in developing norms that preserve the integrity of the community. Reviewers will be specifically instructed to not penalize honesty concerning limitations.
3. **Theory assumptions and proofs** Question: For each theoretical result, does the paper provide the full set of assumptions and a complete (and correct) proof? Answer: [Yes] Justification: To the best of our knowledge, our findings are purely empirical in nature. Guidelines:
  - The answer [N/A] means that the paper does not include theoretical results.
  - All the theorems, formulas, and proofs in the paper should be numbered and crossreferenced.
  - All assumptions should be clearly stated or referenced in the statement of any theorems.
  - The proofs can either appear in the main paper or the supplemental material, but if they appear in the supplemental material, the authors are encouraged to provide a short proof sketch to provide intuition.
  - Inversely, any informal proof provided in the core of the paper should be complemented by formal proofs provided in appendix or supplemental material.
  - Theorems and Lemmas that the proof relies upon should be properly referenced.
4. **Experimental result reproducibility** Question: Does the paper fully disclose all the information needed to reproduce the main experimental results of the paper to the extent that it affects the main claims and/or conclusions of the paper (regardless of whether the code and data are provided or not)? Answer: [Yes] Justification: Please find that the Methods section provides a list of methods, the supplementary section provides the compute resources estimated. Guidelines:
  - The answer [N/A] means that the paper does not include experiments.
  - If the paper includes experiments, a [No] answer to this question will not be perceived well by the reviewers: Making the paper reproducible is important, regardless of whether the code and data are provided or not.
  - If the contribution is a dataset and/or model, the authors should describe the steps taken to make their results reproducible or verifiable.
  - Depending on the contribution, reproducibility can be accomplished in various ways. For example, if the contribution is a novel architecture, describing the architecture fully might suffice, or if the contribution is a specific model and empirical evaluation, it may be necessary to either make it possible for others to replicate the model with the same dataset, or provide access to the model. In general. releasing code and data is often one good way to accomplish this, but reproducibility can also be provided via detailed instructions for how to replicate the results, access to a hosted model (e.g., in the case of a large language model), releasing of a model checkpoint, or other means that are appropriate to the research performed.
  - While NeurIPS does not require releasing code, the conference does require all submissions to provide some reasonable avenue for reproducibility, which may depend on the nature of the contribution. For example
    a. If the contribution is primarily a new algorithm, the paper should make it clear how to reproduce that algorithm.
    b. If the contribution is primarily a new model architecture, the paper should describe the architecture clearly and fully.
    c. If the contribution is a new model (e.g., a large language model), then there should either be a way to access this model for reproducing the results or a way to reproduce the model (e.g., with an open-source dataset or instructions for how to construct the dataset).
    d. We recognize that reproducibility may be tricky in some cases, in which case authors are welcome to describe the particular way they provide for reproducibility. In the case of closed-source models, it may be that access to the model is limited in some way (e.g., to registered users), but it should be possible for other researchers to have some path to reproducing or verifying the results.
5. **Open access to data and code** Question: Does the paper provide open access to the data and code, with sufficient instructions to faithfully reproduce the main experimental results, as described in supplemental material? Answer: [Yes] Justification: Please note that the Dataset used in this paper is a public dataset which belongs to a third party.The code is attached in the zip file alongside the supplementary material. Guidelines:
  - The answer [N/A] means that paper does not include experiments requiring code.
  - Please see the NeurIPS code and data submission guidelines (https://neurips.cc/public/guides/CodeSubmissionPolicy) for more details.
  - While we encourage the release of code and data, we understand that this might not be possible, so [No] is an acceptable answer. Papers cannot be rejected simply for not including code, unless this is central to the contribution (e.g., for a new open-source benchmark).
  - The instructions should contain the exact command and environment needed to run to reproduce the results. See the NeurIPS code and data submission guidelines (https://neurips.cc/public/guides/CodeSubmissionPolicy) for more details.
  - The authors should provide instructions on data access and preparation, including how to access the raw data, preprocessed data, intermediate data, and generated data, etc.
  - The authors should provide scripts to reproduce all experimental results for the new proposed method and baselines. If only a subset of experiments are reproducible, they should state which ones are omitted from the script and why.
  - At submission time, to preserve anonymity, the authors should release anonymized versions (if applicable).
  - Providing as much information as possible in supplemental material (appended to the paper) is recommended, but including URLs to data and code is permitted.
6. **Experimental setting/details** Question: Does the paper specify all the training and test details (e.g., data splits, hyperparameters, how they were chosen, type of optimizer) necessary to understand the results? Answer: [Yes] Justification: Please find the above in the Methods section. Guidelines:
  - The answer [N/A] means that the paper does not include experiments.
  - The experimental setting should be presented in the core of the paper to a level of detail that is necessary to appreciate the results and make sense of them.
  - The full details can be provided either with the code, in appendix, or as supplemental material.
7. **Experiment statistical significance** Question: Does the paper report error bars suitably and correctly defined or other appropriate information about the statistical significance of the experiments? Answer: [Yes] Justification: Please find that we have performed a Cluster mass significance test and omnibus test, large sample set of 25000 samples. Guidelines:
  - The answer [N/A] means that the paper does not include experiments.
  - The authors should answer [Yes] if the results are accompanied by error bars, confidence intervals, or statistical significance tests, at least for the experiments that support the main claims of the paper.
  - The factors of variability that the error bars are capturing should be clearly stated (for example, train/test split, initialization, random drawing of some parameter, or overall run with given experimental conditions).
  - The method for calculating the error bars should be explained (closed form formula, call to a library function, bootstrap, etc.)
  - The assumptions made should be given (e.g., Normally distributed errors).
  - It should be clear whether the error bar is the standard deviation or the standard error of the mean.
  - It is OK to report 1-sigma error bars, but one should state it. The authors should preferably report a 2-sigma error bar than state that they have a 96% CI, if the hypothesis of Normality of errors is not verified.
  - For asymmetric distributions, the authors should be careful not to show in tables or figures symmetric error bars that would yield results that are out of range (e.g., negative error rates).
  - If error bars are reported in tables or plots, the authors should explain in the text how they were calculated and reference the corresponding figures or tables in the text.
8. **Experiments compute resources** Question: For each experiment, does the paper provide sufficient information on the computer resources (type of compute workers, memory, time of execution) needed to reproduce the experiments? Answer: [Yes] Justification: Please find the above in the Supplementary section Guidelines:
  - The answer [N/A] means that the paper does not include experiments.
  - The paper should indicate the type of compute workers CPU or GPU, internal cluster, or cloud provider, including relevant memory and storage.
  - The paper should provide the amount of compute required for each of the individual experimental runs as well as estimate the total compute.
  - The paper should disclose whether the full research project required more compute than the experiments reported in the paper (e.g., preliminary or failed experiments that didn’t make it into the paper).
9. **Code of ethics** Question: Does the research conducted in the paper conform, in every respect, with the NeurIPS Code of Ethics https://neurips.cc/public/EthicsGuidelines? Answer: [Yes] Justification: We have read the code of ethics and abide by it. Guidelines:
  - The answer [N/A] means that the authors have not reviewed the NeurIPS Code of Ethics.
  - If the authors answer [No], they should explain the special circumstances that require a deviation from the Code of Ethics.
  - The authors should make sure to preserve anonymity (e.g., if there is a special consideration due to laws or regulations in their jurisdiction).
10. **Broader impacts** Question: Does the paper discuss both potential positive societal impacts and negative societal impacts of the work performed? Answer: [N/A] Justification: foundational basic science Guidelines:
  - The answer [N/A] means that there is no societal impact of the work performed.
  - If the authors answer [N/A] or [No], they should explain why their work has no societal impact or why the paper does not address societal impact.
  - Examples of negative societal impacts include potential malicious or unintended uses (e.g., disinformation, generating fake profiles, surveillance), fairness considerations (e.g., deployment of technologies that could make decisions that unfairly impact specific groups), privacy considerations, and security considerations.
  - The conference expects that many papers will be foundational research and not tied to particular applications, let alone deployments. However, if there is a direct path to any negative applications, the authors should point it out. For example, it is legitimate to point out that an improvement in the quality of generative models could be used to generate Deepfakes for disinformation. On the other hand, it is not needed to point out that a generic algorithm for optimizing neural networks could enable people to train models that generate Deepfakes faster.
  - The authors should consider possible harms that could arise when the technology is being used as intended and functioning correctly, harms that could arise when the technology is being used as intended but gives incorrect results, and harms following from (intentional or unintentional) misuse of the technology.
  - If there are negative societal impacts, the authors could also discuss possible mitigation strategies (e.g., gated release of models, providing defenses in addition to attacks, mechanisms for monitoring misuse, mechanisms to monitor how a system learns from feedback over time, improving the efficiency and accessibility of ML).
11. **Safeguards** Question: Does the paper describe safeguards that have been put in place for responsible release of data or models that have a high risk for misuse (e.g., pre-trained language models, image generators, or scraped datasets)? Answer: [N/A] Justification: none Guidelines:
  - The answer [N/A] means that the paper poses no such risks.
  - Released models that have a high risk for misuse or dual-use should be released with necessary safeguards to allow for controlled use of the model, for example by requiring that users adhere to usage guidelines or restrictions to access the model or implementing safety filters.
  - Datasets that have been scraped from the Internet could pose safety risks. The authors should describe how they avoided releasing unsafe images.
  - We recognize that providing effective safeguards is challenging, and many papers do not require this, but we encourage authors to take this into account and make a best faith effort.
12. **Licenses for existing assets** Question: Are the creators or original owners of assets (e.g., code, data, models), used in the paper, properly credited and are the license and terms of use explicitly mentioned and properly respected? Answer: [Yes] Justification: Please find that the the origin of the dataset and methods have been clearly cited in the main paper. Guidelines:
  - The answer [N/A] means that the paper does not use existing assets.
  - The authors should cite the original paper that produced the code package or dataset.
  - The authors should state which version of the asset is used and, if possible, include a URL.
  - The name of the license (e.g., CC-BY 4.0) should be included for each asset.
  - For scraped data from a particular source (e.g., website), the copyright and terms of service of that source should be provided.
  - If assets are released, the license, copyright information, and terms of use in the package should be provided. For popular datasets, paperswithcode.com/datasets has curated licenses for some datasets. Their licensing guide can help determine the license of a dataset.
  - For existing datasets that are re-packaged, both the original license and the license of the derived asset (if it has changed) should be provided.
  - If this information is not available online, the authors are encouraged to reach out to the asset’s creators.
13. **New assets** Question: Are new assets introduced in the paper well documented and is the documentation provided alongside the assets? Answer: [N/A] Justification:Please find that only existing methods have been implemented, claims are empirical. Guidelines:
  - The answer [N/A] means that the paper does not release new assets.
  - Researchers should communicate the details of the dataset/code/model as part of their submissions via structured templates. This includes details about training, license, limitations, etc.
  - The paper should discuss whether and how consent was obtained from people whose asset is used.
  - At submission time, remember to anonymize your assets (if applicable). You can either create an anonymized URL or include an anonymized zip file.
14. **Crowdsourcing and research with human subjects** Question: For crowdsourcing experiments and research with human subjects, does the paper include the full text of instructions given to participants and screenshots, if applicable, as well as details about compensation (if any)? Answer: [N/A] Justification: Please note that we did not use crowdsourcing or human subjects. Guidelines:
  - The answer [N/A] means that the paper does not involve crowdsourcing nor research with human subjects.
  - Including this information in the supplemental material is fine, but if the main contribution of the paper involves human subjects, then as much detail as possible should be included in the main paper.
  - According to the NeurIPS Code of Ethics, workers involved in data collection, curation, or other labor should be paid at least the minimum wage in the country of the data collector.
15. **Institutional review board (IRB) approvals or equivalent for research with human subjects** Question: Does the paper describe potential risks incurred by study participants, whether such risks were disclosed to the subjects, and whether Institutional Review Board (IRB) approvals (or an equivalent approval/review based on the requirements of your country or institution) were obtained? Answer: [N/A] Justification:none Guidelines:
  - The answer [N/A] means that the paper does not involve crowdsourcing nor research with human subjects.
  - Depending on the country in which research is conducted, IRB approval (or equivalent) may be required for any human subjects research. If you obtained IRB approval, you should clearly state this in the paper.
  - We recognize that the procedures for this may vary significantly between institutions and locations, and we expect authors to adhere to the NeurIPS Code of Ethics and the guidelines for their institution.
  - For initial submissions, do not include any information that would break anonymity (if applicable), such as the institution conducting the review.
16. **Declaration of LLM usage** Question: Does the paper describe the usage of LLMs if it is an important, original, or non-standard component of the core methods in this research? Note that if the LLM is used only for writing, editing, or formatting purposes and does *not* impact the core methodology, scientific rigor, or originality of the research, declaration is not required. Answer: [Yes] Justification:Drafting sections of the paper, Editing (e.g., grammar, spelling, word choice), Implementing standard methods Guidelines:
  - The answer [N/A] means that the core method development in this research does not involve LLMs as any important, original, or non-standard components.
  - Please refer to our LLM policy in the NeurIPS handbook for what should or should not be described.

## Notes

### Competing Interest Statement

The authors have declared no competing interest.

