## Supplemental Material for "Inferotemporal Cortex Joins the Circuit Before the Code: Non-Serial Inter-Area Synergy in the Macaque Ventral Stream"

### Supplementary Material

**Arun Ram Ponnambalam**
Department of Biomedical Engineering
SRM Institute of Science and Technology

**Krishnan Venkiteswaran Pottore**
Department of Computer Science
University College Dublin

##### Gaussian Copula Mutual Information PID

To decompose information carried by visual cortical areas V1, IT, and V4 about stimulus identity, we applied Partial Information Decomposition using a Gaussian Copula Mutual Information (GCMI) estimator.

##### Copula estimator

Raw neural responses in each ROI were first reduced via MP-PCA (as in the main paper). Rather than applying the Gaussian plug-in estimator directly to the resulting components—which violates Gaussianity assumptions for neural data—we applied a rank-based normalisation to each marginal prior to MI estimation. Specifically, for a variable $\mathbf{X} \in \mathbb{R}^{n \times d}$, each column was transformed via the probit function:

$$\tilde{X}_{ij} = \Phi^{-1}\left(\frac{r_{ij} - 0.5}{n}\right), \tag{1}$$

where $r_{ij}$ is the rank of $X_{ij}$ within column $j$ and $\Phi^{-1}$ is the standard normal quantile function. This marginal Gaussianisation preserves the rank-order dependence structure while rendering the Gaussian MI formula applicable. MI was then estimated from covariances of the transformed data:

$$\widehat{I}(X;Y) = \frac{1}{2}\left(\log|\Sigma_X| + \log|\Sigma_Y| - \log|\Sigma_{XY}|\right), \tag{2}$$

where $\Sigma_X$, $\Sigma_Y$, and $\Sigma_{XY}$ are the covariance matrices of $\tilde{\mathbf{X}}$, $\mathbf{Y}$, and their concatenation, respectively. Only the neural component $\mathbf{X}$ was Gaussianised; CNN feature targets $\mathbf{Y}$ (PCA-50 of ResNet-18 avgpool activations) were used as-is.

##### PID decomposition

Total MI from the three ROIs jointly was decomposed into redundancy, unique, and synergistic atoms following the minimum-MI redundancy axiom:

$$\text{Red} = \min_{s \in \{V1,\, IT,\, V4\}} \widehat{I}(S_s; Y), \tag{3}$$

$$\text{Uniq}_s = \max\Big(0,\ \widehat{I}(S_s; Y) - \text{Red}\Big), \tag{4}$$

$$\text{Syn} = \max\Big(0,\ \widehat{I}(Z_{\text{joint}}; Y) - \text{Red} - \textstyle\sum_s \text{Uniq}_s\Big). \tag{5}$$

#### 1 Computational Resources

All experiments and analyses were conducted using Google Colaboratory (Colab). Computations were performed utilizing an NVIDIA A100 Tensor Core GPU with the High-RAM runtime configuration.

Preprint.

Table 1: **Supplementary Table S1:** Participation Ratio (PR) Quench Magnitudes and Temporal Latencies. Baseline PR is calculated as the mean dimensionality from 0–100 ms. Quench Time indicates the latency of the post-stimulus dimensionality minimum.

| | Monkey N | | | | Monkey F | | | |
|---|---|---|---|---|---|---|---|---|
| **ROI** | **Baseline** (PR) | **Min PR** (PR) | **Time** (ms) | **Quench %** ($\Delta$%) | **Baseline** (PR) | **Min PR** (PR) | **Time** (ms) | **Quench %** ($\Delta$%) |
| V1 | 34.47 | 11.20 | 142.5 | 67.51% | 27.21 | 21.56 | 132.5 | 20.76% |
| V4 | 28.96 | 16.74 | 142.5 | 42.20% | 27.03 | 25.88 | 147.5 | 4.26% |
| IT | 24.57 | 18.16 | 192.5 | 26.10% | 14.24 | 12.65 | 152.5 | 11.17% |

Table 2: **Supplementary Table S2:** Peak Informational Metrics across the Ventral Stream Hierarchy. $I(S; \mathbf{R})$ represents the total joint mutual information between the three cortical areas (V1, V4, IT) and the specific CNN layer representation. All values are reported in nats.

| | Monkey N | | | Monkey F | | |
|---|---|---|---|---|---|---|
| **CNN Target Layer** | **Peak** $I(S; \mathbf{R})$ | **Peak** $Syn$ | **Peak** $Red$ | **Peak** $I(S; \mathbf{R})$ | **Peak** $Syn$ | **Peak** $Red$ |
| ResNet18-Layer 1 | 3.25 | 0.56 | 0.97 | 2.85 | 1.02 | 1.09 |
| ResNet18-Layer 2 | 3.34 | 0.94 | 1.24 | 3.17 | 1.42 | 1.38 |
| ResNet18-Layer 3 | 3.42 | 1.34 | 1.49 | 3.43 | 1.56 | 1.43 |
| ResNet18-Layer 4 | 1.94 | 0.60 | 0.66 | 1.97 | 0.71 | 0.55 |
| ResNet18-AvgPool | 1.94 | 0.60 | 0.66 | 1.97 | 0.71 | 0.55 |

This hardware environment provided the necessary memory capacity and compute capability to efficiently execute the models and process the neural population datasets. The source code required to reproduce all experimental results is included in our supplementary code submission.

Table 3: **Supplementary Table S3:** Comparative Onset Latencies of Synergy and Unique IT Information. The $\Delta$ (ms) represents the temporal dissociation between circuit-level integrated activity and the settlement of independent IT representations.

| | Monkey N | | | Monkey F | | |
|---|---|---|---|---|---|---|
| **CNN Target Layer** | **Syn Onset** (ms) | **IT Onset** (ms) | $\Delta$ (ms) | **Syn Onset** (ms) | **IT Onset** (ms) | $\Delta$ (ms) |
| ResNet18-Layer 1 | 127.5 | — | — | 162.5 | 177.5 | 15.0 |
| ResNet18-Layer 2 | 127.5 | — | — | 162.5 | 177.5 | 15.0 |
| ResNet18-Layer 3 | 127.5 | 197.5 | 70.0 | 122.5 | 172.5 | 50.0 |
| ResNet18-Layer 4 | 132.5 | 182.5 | 50.0 | 137.5 | 167.5 | 30.0 |
| ResNet18-AvgPool | 132.5 | 182.5 | 50.0 | 137.5 | 167.5 | 30.0 |

Table 4: **Supplementary Table S4:** Comprehensive Peak PID Metrics across the ResNet-18 Hierarchy for Monkey N and Monkey F. All values are reported in nats.

| | Monkey N | | | | | | Monkey F | | | | | |
|---|---|---|---|---|---|---|---|---|---|---|---|---|
| **Layer** | **Red** | $U_{V1}$ | $U_{V4}$ | $U_{IT}$ | **Syn** | $I_{joint}$ | **Red** | $U_{V1}$ | $U_{V4}$ | $U_{IT}$ | **Syn** | $I_{joint}$ |
| Layer 1 | 0.97 | 1.61 | 1.22 | 0.00 | 0.56 | 3.25 | 1.09 | 2.01 | 1.05 | 0.18 | 1.02 | 2.85 |
| Layer 2 | 1.24 | 1.34 | 1.37 | 0.00 | 0.94 | 3.34 | 1.38 | 1.75 | 1.07 | 0.36 | 1.42 | 3.17 |
| Layer 3 | 1.49 | 1.04 | 1.32 | 0.07 | 1.34 | 3.42 | 1.43 | 1.54 | 1.00 | 0.78 | 1.56 | 3.43 |
| Layer 4 | 0.66 | 0.39 | 0.70 | 0.31 | 0.60 | 1.94 | 0.55 | 0.62 | 0.54 | 1.02 | 0.71 | 1.97 |
| AvgPool | 0.66 | 0.39 | 0.70 | 0.31 | 0.60 | 1.94 | 0.55 | 0.62 | 0.54 | 1.02 | 0.71 | 1.97 |

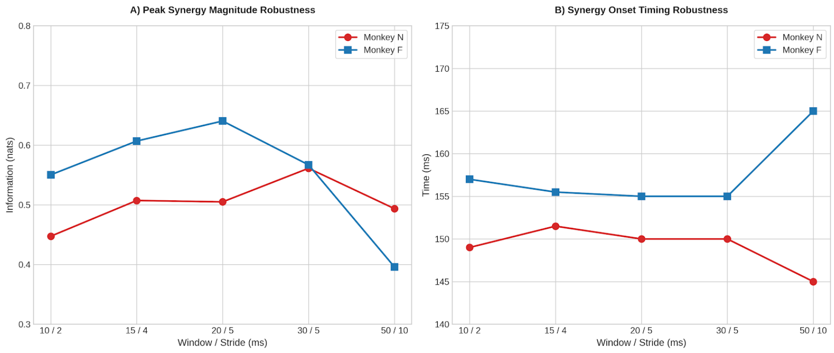


Figure 1: **(a) Stability of peak synergistic magnitude.** The maximum synergistic information (nats) remains robustly between 0.4 and 0.7 nats across both subjects (Monkey N, red circles; Monkey F, blue squares) for window lengths ranging from 10 ms to 50 ms. This invariance indicates that the synergistic component of the neural population code is a stable feature of the circuit's information topology, rather than an artifact of a specific integration timescale. **(b) Invariance of synergy onset timing.** The temporal onset of synergistic circuit activity (ms post-stimulus) is tightly constrained between 145 ms and 165 ms across the entire parameter sweep. This stability validates that the "synergy-first" principle is an intrinsic property of the $V1 \rightarrow V4 \rightarrow IT$ interaction, remaining independent of the sliding-window configuration used for the 3-source Partial Information Decomposition (PID) analysis.

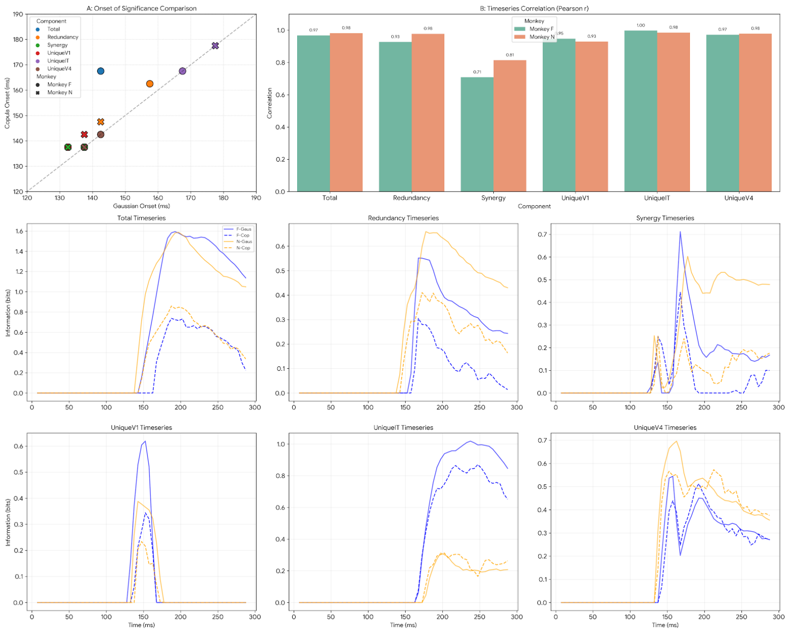


Figure 2: Validation of Gaussian Plug-in vs. Copula PID Estimators. Comprehensive comparison of the 3-source PID components across all monkeys. **(A) Onset of Significance Comparison.** Scatter plot illustrating high temporal consistency in the detection of information emergence across all PID atoms (Total MI, Redundancy, Synergy, and Unique information for V1, V4, and IT). **(B) Timeseries Correlation.** Pearson correlation coefficients ($r$) between the Gaussian and Copula timeseries for each PID component. High correlation values (typically $r > 0.80$) demonstrate that the Gaussian plug-in effectively captures the underlying neural population dynamics. **(Bottom Panels) Component Timeseries.** Representative temporal trajectories for each PID atom. Solid lines represent Gaussian plug-in estimates and dashed lines represent Copula estimates (Blue: Monkey F, Orange: Monkey N), highlighting the matched temporal envelopes across both monkeys and all estimators.

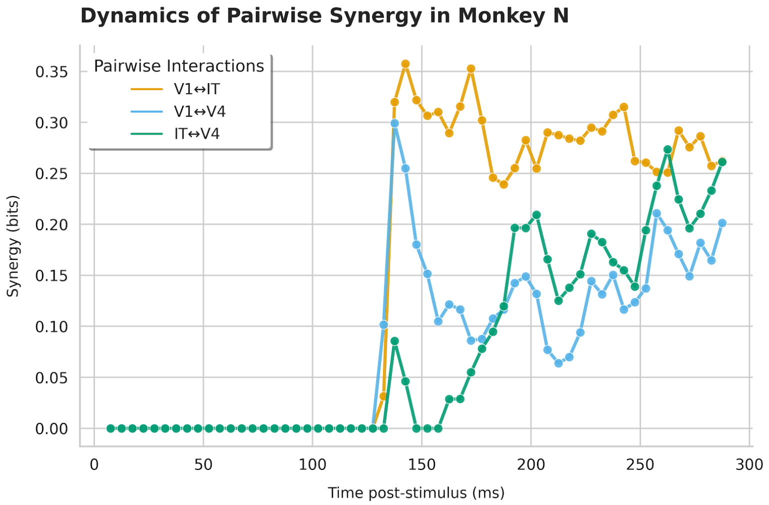


Figure 3: **Dynamics of Pairwise Synergy using GCMI**

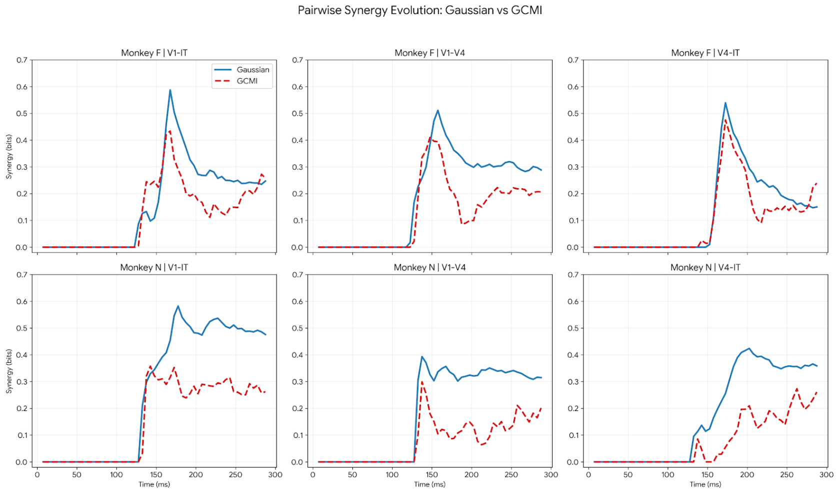


Figure 4: **Pairwise Synergy Evolution: Gaussian vs. GCMI.** Temporal dynamics of synergy calculated for cortical pairings (V1-IT, V1-V4, and V4-IT) across two subjects (Monkey F, top; Monkey N, bottom). Solid blue lines represent the Gaussian plug-in estimator (derived from spatially pooled Average Pool layer activations), while dashed red lines denote the GCMI (Copula-based) estimator. The plots demonstrate that while the Gaussian estimator yields higher absolute bit values, both estimators exhibit highly synchronized temporal profiles, identifying identical peaks and fluctuations in synergy across all area pairs and both monkeys.

Table 5: **Supplementary Table S5:** Significant Onset Latencies for Pairwise Synergy across the ResNet-18 Hierarchy. Values represent ms post-stimulus. Threshold for significance is $\geq 0.05$ nats.

| | Monkey N | | | Monkey F | | |
|---|---|---|---|---|---|---|
| **Layer** | $V1 \leftrightarrow IT$ | $V1 \leftrightarrow V4$ | $IT \leftrightarrow V4$ | $V1 \leftrightarrow IT$ | $V1 \leftrightarrow V4$ | $IT \leftrightarrow V4$ |
| ResNet18-Layer 1 | 127.5 | 127.5 | 132.5 | 122.5 | 117.5 | 157.5 |
| ResNet18-Layer 2 | 127.5 | 127.5 | 132.5 | 122.5 | 117.5 | 157.5 |
| ResNet18-Layer 3 | 127.5 | 127.5 | 132.5 | 122.5 | 122.5 | 157.5 |
| ResNet18-Layer 4 | 132.5 | 132.5 | 132.5 | 127.5 | 127.5 | 157.5 |
| ResNet18-AvgPool | 132.5 | 132.5 | 132.5 | 127.5 | 127.5 | 157.5 |
